# Chronic Exposure to Glucocorticoids Induces Suboptimal Decision-Making in Mice

**DOI:** 10.1101/2020.09.11.293217

**Authors:** Lidia Cabeza, Bahrie Ramadan, Julie Giustiniani, Christophe Houdayer, Yann Pellequer, Damien Gabriel, Sylvie Fauconnet, Emmanuel Haffen, Pierre-Yves Risold, Dominique Fellmann, David Belin, Yvan Peterschmitt

## Abstract

Anxio-depressive symptoms as well as severe cognitive dysfunction including aberrant decision-making (DM) are documented in neuropsychiatric patients with hypercortisolaemia. Yet, the influence of the hypothalamo-pituitary-adrenal (HPA) axis on DM processes remains poorly understood. As a tractable mean to approach this human condition, adult male C57BL/6JRj mice were chronically treated with corticosterone (CORT) prior to behavioural, physiological and neurobiological evaluation. The behavioural data indicate that chronic CORT delays the acquisition of contingencies required to orient responding towards optimal DM performance in a mouse Gambling Task (mGT). Specifically, CORT-treated animals show a longer exploration and a delayed onset of the optimal DM performance. Remarkably, the proportion of individuals performing suboptimally in the mGT is increased in the CORT condition. This variability seems to be better accounted for by variations in sensitivity to negative rather than to positive outcome. Besides, CORT-treated animals perform worse than control animals in a spatial working memory (WM) paradigm and in a motor learning task. Finally, Western blotting neurobiological analyses show that chronic CORT downregulates glucocorticoid receptor expression in the medial Prefrontal Cortex (mPFC). Besides, corticotropin-releasing factor signalling in the mPFC of CORT individuals negatively correlates with their DM performance. Collectively, this study describes how chronic exposure to glucocorticoids induces suboptimal DM under uncertainty in a mGT, hampers WM and motor learning processes, thus affecting specific emotional, motor, cognitive and neurobiological endophenotypic dimensions relevant for precision medicine in biological psychiatry.

## INTRODUCTION

Chronically elevated circulating glucocorticoids (GC) have been extensively shown to have detrimental physiological and cognitive effects (see for instance [1]). Particularly, persistent hypothalamo-pituitary-adrenal (HPA) axis dysfunction has been reported in humans upon repeated stress, with elevated levels of the endogenous GC cortisol [2–4], but also in patients with chronic inflammatory diseases treated with exogenous GC [5–7]. In fact, hypercortisolaemia is part of the symptomatology reported in patients with neuropsychiatric disorders afflicted with severe cognitive dysfunction [8,9]. Specifically, aberrant decision-making (DM) has been described in patients suffering from depression using the Iowa Gambling Task (IGT) [10]. This paradigm involves probabilistic learning via monetary rewards and penalties, and optimal task performance that leads to the maximization of gains, requires subjects to develop a preference for smaller immediate rewards in order to avoid more important losses in the long-term. Interestingly, maladaptive DM strategies have also been reported in healthy subjects [11,12]. Of particular interest, depressed patients show a reduced ability to detect and incorporate experience from reward-learning associations [13], therefore anhedonia is thought to act by modifying goal-directed behaviours when positive reinforcements are involved [14]. Moreover, hyposensitivity to positive outcome (reward) and maladaptive responses to negative outcome have been linked to depression [15,16], suggesting a dysfunctional interaction between limbic and motor-executive regions as putative underlying mechanisms. Yet, the influence of the HPA axis on DM alterations remains poorly understood. The regulatory role of GC on HPA axis activity [17] has pointed to imbalances in the expression of their main receptors (glucocorticoid-GR, and mineralocorticoid receptors – MR) as biomarkers of depressive states [18,19]. Simultaneously, the corticotropin-releasing factor (CRF), the major activator of the HPA axis, is thought a key player in stress-induced executive dysfunction [20] and in mood and anxiety disorders [21–25].

Chronic corticosterone (CORT) administration in rodents represents a tractable mean to address these human pathological conditions [26]. In fact, chronic CORT-treated animals exhibit a behavioural spectrum reminiscent to emotional anxio-depressive symptoms as evidenced in several non-conditioned tasks [27–29]. Besides, in gambling tasks, healthy rodents efficiently explore and sample from different options prior to establish their choice strategy upon associative and reinforcement learning, showing a high inter-individual variability, probably shared with humans [30–36].

Here, we hypothesized that chronic CORT exposure leads to suboptimal DM processing under uncertainty in a mouse Gambling Task. In line with the dimensional framework of the Research Domain Criteria Initiative (RDoC) [37], we addressed feedback sensitivity since optimal performance in gambling tasks requires effective exploration of options in their early stages [32,33]. Aiming to elucidate their implication in suboptimal DM, spatial working memory (WM) and psychomotricity, as cognitive and arousal-sensorimotor constructs, were also explored. Three relevant brain regions were targeted in this study given their contribution in instrumental behaviour, and mood and stress-related symptomatology: the medial prefrontal cortex (mPFC) and the dorsolateral striatum (DLS), modulators of goal-directed and habit-based learning processes respectively [38,39], and the ventral hippocampus (VH), involved in stress and emotional processing, exerting strong regulatory control on the HPA axis [40]. The protein levels of GR, MR and CRF were quantified in these brain areas. As depression is associated with a high rate of pharmacological resistance [41,42] and to a high risk of suicide [43,44], understanding how neuronal mechanisms underlying DM processes are altered may offer insights towards the detection of predictive biomarkers for treatment selection.

## MATERIALS AND METHODS

### Animals

Eighty 6-8 week-old male C57BL/6JRj mice (*EtsJanvier Labs*, Saint-Berthevin, France) were group-housed and maintained under a normal 12-hour light/dark cycle with constant temperature (22±2°C). They had access to standard chow (Kliba Nafag 3430PMS10, *Serlab*, CH-4303 Kaiserau, Germany) *ad libitum* for three weeks, and the fourth week onwards, under food restriction to 80-90% of their free-feeding weight (mean ± SEM (g) = 26.20±0.26). Bottles containing water and/or treatment were available at all times.

Experiments were all conducted following the standards of the Ethical Committee in Animal Experimentation from Besançon (CEBEA-58; A-25-056-2). All efforts were made to minimize animal suffering during the experiments according to the Directive from the European Council at 22^nd^ of September 2010 (2010/63/EU).

### Pharmacological treatment

Mice started being treated four weeks before the beginning of the behavioural assessment. Half the individuals received corticosterone (CORT, -4-Pregnene-11β-diol-3,20-dione-21-dione, *Sigma-Aldrich*, France) in the drinking water (35μg/ml equivalent to 5 mg/kg/day, CORT group). CORT was freshly dissolved twice a week in vehicle (0.45% hydroxypropyl-β-cyclodextrin-βCD, *Roquette GmbH*, France) which control animals (SHAM group) received in the drinking water throughout the entire experiment.

### Multi-domain behavioural characterization

Mice were tested behaviourally during the light phase of the cycle (from 8:00 a.m.) after 4 weeks of differential treatment. A timeline of the experiment is presented in **Figure 1**, established within the framework of the RDoC to assess the functioning of several complementary systems, including Negative and Positive Valence Systems, Cognitive, Sensorimotor and Arousal and Regulation Systems.

**Figure 1.**
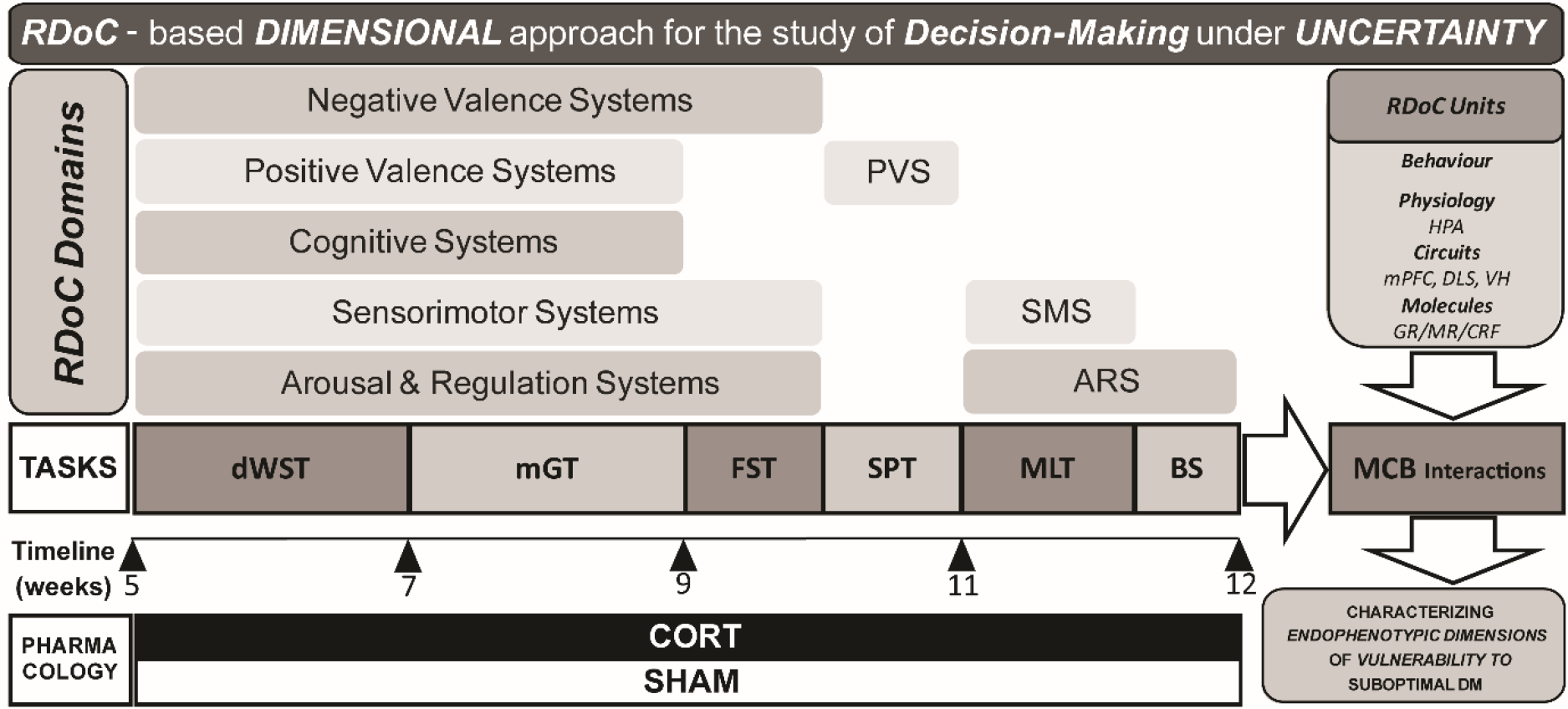
Dimensional approach of the experimental design. Mice were treated with corticosterone (CORT) (35μg/mL; n=40) or vehicle (0.45% hydroxypropil-β-cyclodextrine-βCD; n=40) for four weeks before behavioural screening started. Treatment was maintained throughout. Mice were tested in a series of behavioural tasks, namely delayed spatial Win Shift Task (dWST), mouse Gambling Task (mGT), Forced Swim Test (FST), Sucrose Preference Test (SPT) and Motor Learning Task (MLT) in order to assess the functioning of several *Research Domain Criteria* (RDoC) domains/systems including Negative and Positive Valence Systems (PVS), Cognitive Systems, Sensorimotor Systems (SMS) and Arousal and Regulation Systems (ARS). Five-seven days after the last behavioural test, mice were subjected to blood sampling (BS) and then sacrificed. Their brains were harvested for subsequent post-mortem analyses enabling a multilevel approach, e.g., molecules–circuits–behaviour (MCB) (Hyphothalamo-pituitairy-adrenal axis –HPA; medial prefrontal cortex –mPFC; dorsolateral striatum –DLS; ventral hippocampus –VH; glucocorticoid receptor –GR; mineralocorticoid receptor -MR; corticotropin-releasing factor – CRF), as biobehavioural basis of the influence of chronic CORT exposure.

#### Delayed spatial Win-Shift Task (dWST)

After 5 consecutive training days, spatial WM was tested in a subset of animals (SHAM, n=22; CORT, n=22) as previously described (adapted from [45], **for details see SOM**).

#### Mouse Gambling Task (mGT)

Decision-making was measured using the mGT task the protocol of which we have previously described [30] and is extensively detailed in the **SOM**. Decision-making performance in the mGT was measured as the percentage of advantageous choices over five 20-trial sessions. Choice strategy based on 4 different behavioural dimensions (stickiness, flexibility, lose-shift and win-stay scores), was assessed in 40-trial blocks as previously described [30,32,33,46]. Performance during the last session was considered for the overall measure of DM performance, as previously described [34,35]. Six mice displaying immediate spatial preference among options (choice proportion different from the expected in absence of spatial preference, thus 25% of choices for each of the 4 available options; X^2^, p<0.05) were discarded from the subsequent mGT analyses.

#### Sucrose Preference Test (SPT)

The individual sensitivity to reward [47] was measured using the preference for a sucrose solution over water, as previously described [30].

#### Forced Swim Test (FST)

Coping strategies in the face of distressing, uncertain conditions were measured in a FST [19,48]. Mice were individually placed for 6 minutes in an inescapable glass cylinder filled with 20 cm of warm water (31.5±0.5°C) and the overall time during which they were immobile was recorded [49]. Two animals were discarded from the analysis due to technical reasons.

#### Motor Learning Task (MLT)

Psychomotricity was measured using a rotarod task (adapted from [50]) and detailed in the **SOM**.

All procedures including food reward (20mg Dustless Precision Pellets^®^ Grain-Based Diet, *PHYMEP s.a.r.L.*, Paris, France) were preceded by a habituation period inside the home cages (**see Figure 1** for the experimental design).

### Physiological responses to chronic CORT treatment

#### Fur Coat State (FCS)

The effects of chronic CORT exposure on self-oriented behaviour were assessed using the weekly measurement of the state of the fur coat of each animal, as previously described [51].

#### CORT plasma assays

Final trunk blood samples were collected from all animals (n=80) 5-7 days after the last behavioural test and directly centrifuged at 2100 g for 15 minutes at 20°C. Serum was collected and stored at −80°C until assayed. Plasma CORT concentration was measured using an immunoassay kit (DetectX Corticosterone Immunoassay kit, *Arbor Assays*, Ann Arbor, Michigan, USA). In order to measure the homeostatic stress reactivity of the HPA axis, blood samples were also collected from a subset of mice (SHAM, n=10; CORT, n=9) following a gentle restraint stress [52] directly before sacrifice.

### Western-blots

Animals were sacrificed by rapid cervical dislocation 5-7 days after the last behavioral test, in the central hours of the light cycle (from 2:00 p.m.). Brains were removed, snap-frozen and stored at −80°C until processed. Bilateral samples from the mPFC (from 2.3 to 1.3 mm anterior to bregma), the DLS (from 1.1 to 0.1 anterior to bregma) and the VH (from 2.8 to 3.8 posterior to bregma) were obtained from 1 mm-thick coronal sections obtained using a cryostat and stored at −80°C.

Samples were processed as described in the **SOM** with anti-GR (mouse; 1:500; sc-393232, *Santa Cruz Biotechnology*), anti-MR (rabbit; 1/1000; ab62532, *Abcam*) or anti-CRF (mouse; 1:250; sc-293187, *Santa Cruz Biotechnology*) primary antibodies and HRP anti-Mouse Ig (goat; 1:5000; *BD Pharmingen*™) or HRP anti-Rabbit (goat; 1:5000; *BD Pharmingen*™) secondary antibodies. Membranes were reprobed with anti-β-actin, which served as a loading control and allows normalization for sample comparison (mouse; 1:1000; sc-47778, *Santa Cruz Biotechnology*).

Western-blot images were acquired either with a Bio-Rad ChemiDoc XRS+ System (Life-Sience, *Bio-Rad*, France) or with autoradiographic films (Hyperfilm ECL, GE *Healthcare*, Velizy-Villacoublay, France). All quantifications were made blind to the experimental conditions using ImageJ software (*National Institutes of Health*, Bethesda, MD, USA) (see **Figure S1**).

Due to technical issues, some samples were not included in the final analyses, so that final samples sizes were: mPFC, GR/MR n=72, CRF n=53; DLS, GR/MR n=70, CRF n=55; VH, GR/MR n=66, CRF n=57.

### Data and statistical analyses

Data are presented as means ± SEM.

Statistical analyses were conducted using STATISTICA 10 (*Statsoft*, Palo Alto, USA) and figures were designed using GraphPad Prism 7 software (*GraphPad Inc*., San Diego, USA).

The sample sizes were identified a priori by statistical power analysis (*G*Power* software, Heinrich Heine Universität, Dusseldorf, Germany) with a repeated measures ANOVA (RM-ANOVA) design including 3 groups (between-subject factor), 5 measurements (within-subject factor) and predicted effect size of 0.14, 1-ß=0.8 and α=0.05. Our animal sample is predicted to yield highly reproducible outcomes with 1-ß>0.8 and α<0.05.

Individuals across pharmacological conditions were clustered in three different groups, namely (1) good, (2) intermediate and (3) poor decision-makers (DMs), with distinct preference for the advantageous options: ≥70% preference, between 70% and 50% preference, and ≤50% preference respectively.

Assumptions for parametric analysis were verified prior to each analysis: normality of distribution with Shapiro-Wilk, homogeneity of variance with Levene’s and sphericity with Mauchly’s tests. Behavioural time-dependent measures assessed during the mGT, the dWST and the MLT were analysed by RM-ANOVA with session (1 to 5) or 40-trial block (beginning or end) as within-subject factors, and treatment (SHAM vs CORT) or clusters (good, intermediate or poor DMs) as between-subject factors. Group’s performance in the mGT was compared to chance level (50% of advantageous choices) using Student t-tests. The degradation of the coat state due to the treatment was analysed by ANOVA with factors being weeks (1 to 13) and treatment (SHAM vs CORT). When datasets did not meet assumptions for parametric analyses, non-parametric analyses i.e. Kruskal-Wallis, Wilcoxon or Mann Whitney U tests, were used. Upon significant main effects, further comparisons were performed with Duncan or Bonferroni corrections.

The assumption of independent and normally distributed distribution of treatment populations within each cluster was tested with Chi-squared tests (X^2^). Dimensional relationships between behavioural markers of DM, as measured as final performance (% of advantageous choices in the last session) variable in the mGT and protein levels in the various brain structures under investigation were analysed using Pearson correlations.

For all analyses, alpha was set at 0.05 and effect sizes are reported as partial *η*^2^ (p*η*^2^).

## RESULTS

At the population level, all mice showed a progressive increase in their performance in the mGT over 5 sessions [main effect of session: F_4,288_=31.8, p<0.0000, p*η*^2^=0.31] but no general difference was found between groups [treatment: F_1,72_=2.4, p>0.05, p*η*^2^=0.03] (**Figure 2A**). However, further analyses revealed that whereas SHAM mice allocated their response preferentially towards advantageous options from session 2 onwards [% advantageous choices vs chance, session 1: t_38_=1.2, p>0.05, sessions 2-5: t_38_>2.5, all ps<0.0000], CORT mice required 20 more trials to improve DM [session 1&2: t_34_<2.7, ps>0.05, sessions 3-5: t_34_>4.8, all ps<0.0000]. Thereby this highlights that chronic CORT lengthens exploration and delays the onset of optimal DM performance.

**Figure 2.**
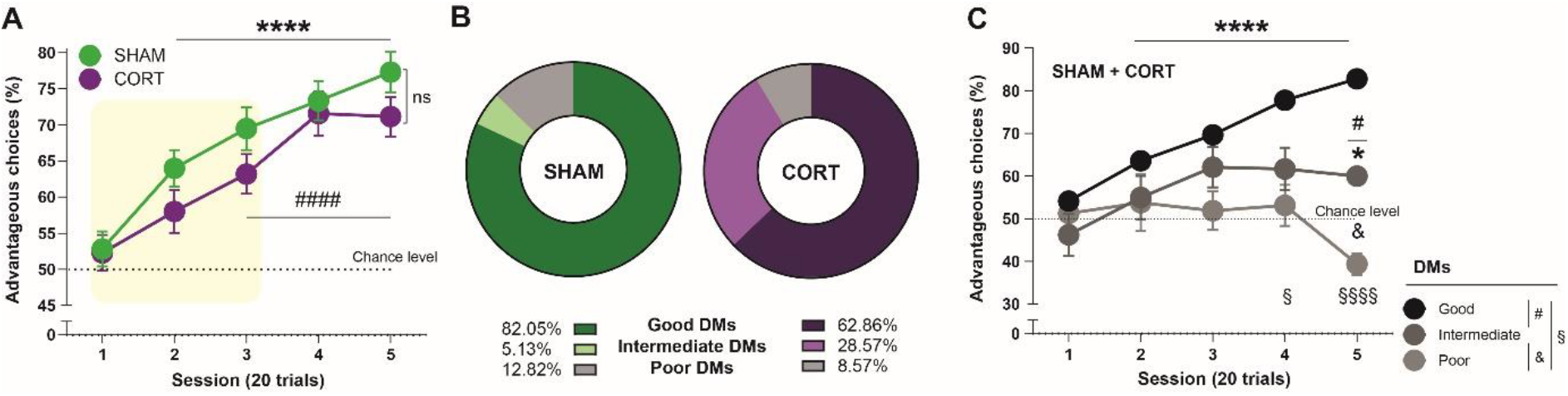
Chronic CORT increases the propensity for suboptimal decision-making in the mouse Gambling Task. (A) The behavioural performance in the mGT of CORT-treated animals reveals a longer exploration of options and a delayed onset of optimal DM strategy compared to controls (SHAM). CORT animals needed 40 trials to allocate their response towards advantageous options, while SHAM animals required only 20 trials [% advantageous choices different from 50%: SHAM, ****, p<0.0000; CORT, ####, p<0.0000]. (B) The frequency distribution of SHAM and CORT individuals in good (n=54), intermediate (n=12) and poor (n=8) DM categories was significantly different [X^2^, p<0.0001], with an increased proportion of treated animals among the intermediate DMs. (C) The three DM subpopulations learnt at different speeds [cluster × session interaction: p<0.0000]. Good (black) DMs improved their performance already in the second session [% advantageous choices different from 50%, ****, p<0.0000], while intermediate (dark grey) DMs only significantly improved in the last session [*, p<0.05]. Good and intermediate DMs strategies significantly differed in the last session [#, p<0.05]. Poor DMs (light grey) however, never allocated their response towards the advantageous options and performed differently than good DMs from the fourth session [session 4: §, p<0.05; session 5: §§§§, p<0.0000], and than intermediate DMs in the last session [&, p<0.05].

We further explored whether chronic CORT could be considered a vulnerabilization factor to suboptimal DM (**Figure 2B**). The majority of individuals displayed the optimal strategy (good DMs). They represent 82.05% of SHAM and only 62.86% of CORT animals (mean % advantageous choices ± SEM: 82.78±1.25). Individuals from the intermediate DM subpopulation developed a delayed preference towards the advantageous options, without reaching the optimal strategy. They constitute only 5.13% of SHAM whereas 28.57% of CORT mice (60.00±1.38). Poor DMs failed to develop a preference for any option and correspond to 12.82% of SHAM and 8.57% of CORT animals (39.38±2.58). The distribution of CORT and SHAM mice in the three subpopulations was compared, highlighting a significant difference (X^2^, p<0.0001) which is mainly accounted for by the intermediate DMs.

In-depth analysis of interindividual variability was further performed. Decision-making subpopulations learnt at different rates [main effect of group: F_2,71_= 21.0, p<0.0000; p*η*^2^=0.37; session × cluster interaction: F_8,284_=6.1, p<0.0000, p*η*^2^=0.15] (**Figure 2C**). Good DMs (n=54) needed 20 trials to orientate towards the advantageous options [% advantageous choices vs chance, session 1: t_53_=2.1, p>0.05, sessions 2-5: t_53_>6.2, all ps<0.0000], while intermediate DMs (n=12) needed 80 trials [session 1-4: Z<2.05, p>0.05, session 5: Z=7.3, p<0.0001]. Unlike the other two categories, poor DMs (n=8) never exhibited a preference [session 1-5: Z<2.4, all ps>0.05]. In the last session of the mGT, good DMs performed differently than intermediate DMs [p<0.05, post-hoc], and the latter differently than poor DMs [p<0.05, post-hoc]. Good and poor DMs performed differently from the fourth session [session 4: p<0.05, session 5: p<0.0001, post-hoc]. Further analyses revealed that, within the good DMs cluster, whose individuals developed the optimal performance, CORT treatment delayed the onset of the strategy i.e. the allocation of the responses towards the advantageous options. Good DMs from the CORT group (n=22) performed differently than chance from the third session onwards [sessions 1&2: Z<2.4, p>0.05, sessions 3-5: Z>3.5, all ps<0.01], while in the SHAM group (n=32) they differed already from the second session [sessions 1: t_31_=1.2, p>0.05, sessions 2-5: t_31_>6.1, all ps<0.0000]. Collectively these data indicate that chronic CORT increases the propensity to suboptimal DM performance with an increased proportion of intermediate DMs compared to controls.

To infer strategies mediating mGT performance, we studied the evolution of the behavioural dimensions along task progression. Stickiness [main effect of block: F_1,72_=54.8, p<0.0000, p*η*^2^=0.43] and flexibility [block: F_1,72_=35.4, p<0.0000, p*η*^2^=0.33] scores changed along the task, without significant effect of the CORT treatment [main effect of treatment, stickiness: F_1,72_=0.6, p>0.05, p*η*^2^=0.01; flexibility: F_1,72_=3.4, p>0.05, p*η*^2^=0.05] (**Figure 3A, 3B**). Final stickiness [r=0.669, p<0.0000] and flexibility scores [r=−0.585, p<0.0000] significantly correlate with final mGT performance.

**Figure 3.**
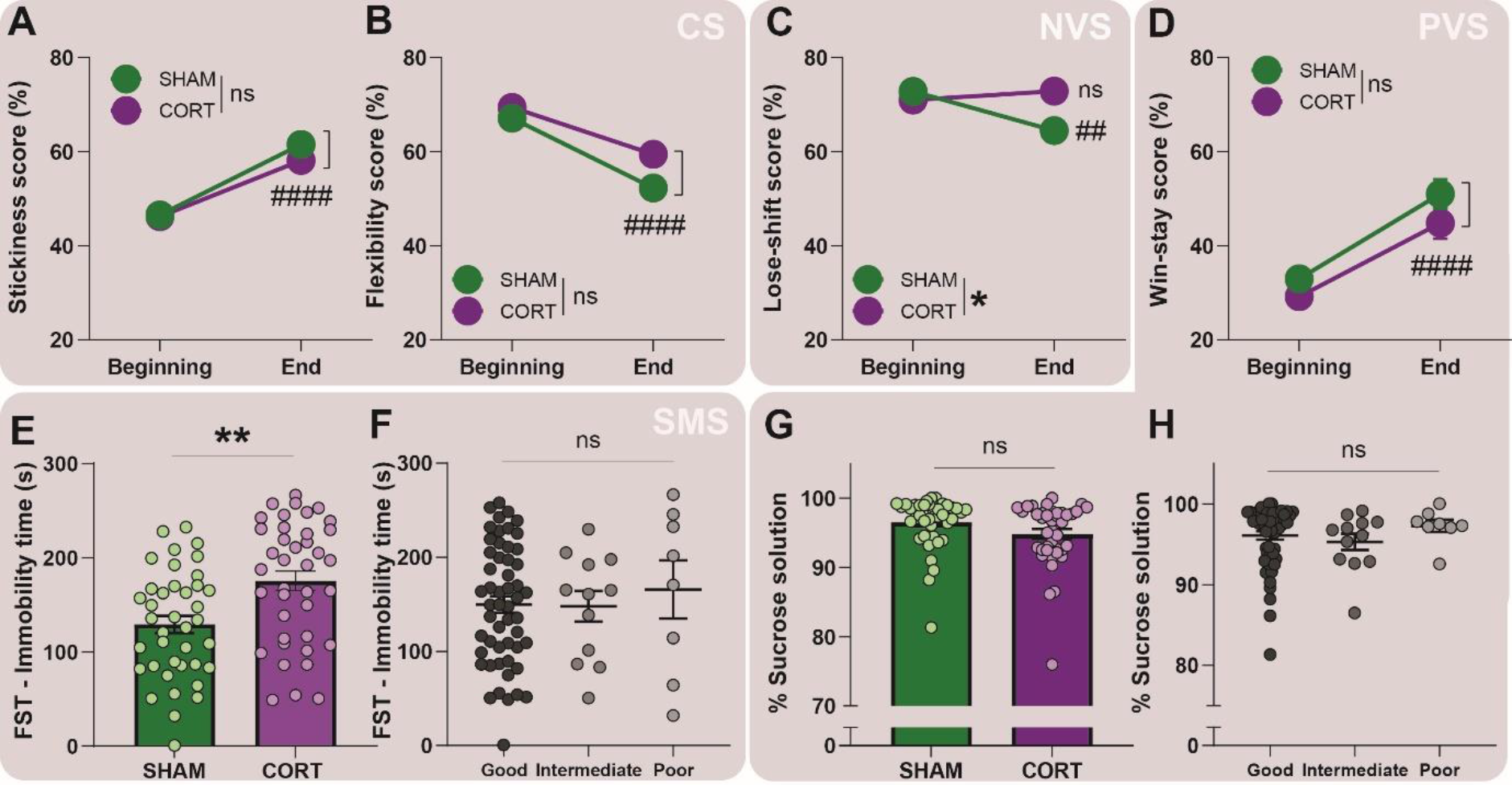
Chronic CORT differentially impacts DM strategy and enhances sensitivity to negative outcome. The evolution of the behavioural dimensions along the mGT was studied in order to evaluate the influence of CORT overexposure in the development of an optimal DM strategy. Irrespective of the treatment [ns], mice became more rigid [####, p<0.0000] (A) and less flexible [####, p<0.0000] (B) in their choices as the task progressed (Cognitive Systems -CS). However, while SHAM animals learnt to cope with penalties [##, p<0.01], CORT animals continued to frequently change option after a negative outcome at the end of the task [ns]. Final lose-shift scores significantly differed between conditions [treatment: *, p<0.05] (Negative Valence Systems -NVS) (C). Moreover, irrespective of the treatment, all animals more frequently selected the same option after a positive outcome with the task progression [####, p<0.0000] (D) (Positive Valence Systems -PVS). Chronic CORT significantly influenced the coping style of mice when facing uncertainty in the Forced Swim Test (FST), with an increment of their immobility duration compared to SHAM animals [treatment: **, p<0.01] (E). However, DM clusters displayed similar coping styles [ns] (F) (Sensorimotor Systems -SMS). In terms of reward sensitivity, no differences were found between SHAM and CORT animals in the preference for a sucrose solution over water [ns] (G), neither between DM subpopulations [ns] (H).

Remarkably, a significant effect of the treatment in interaction with the time course for the lose-shift score was evidenced [block × treatment interaction: F_1,72_=4.7, p<0.05; p*η*^2^=0.06). At the beginning of the task, all animals were prone to option change after a negative outcome (mean percentage ± SEM of lose-shift, SHAM: 72.83±1.89; CORT: 70.97±2.63). At the end of the task, SHAM animals were significantly less prone to change after a penalty (64.49±2.59) than CORT animals (72.85±2.75) [p<0.01, post-hoc] (**Figure 3C**). Whereas lose-shift scores do not correlate with final mGT performance in CORT animals [r=−0.050, p=0.77], a trend was evidenced for the SHAM condition [r=−0.307, p=0.058]. Concerning the win-stay score, all animals more frequently chose the same option after a reward as the task progressed [main effect of blocks: F_1,72_=52.8, p<0.0000, p*η*^2^=0.42], irrespective of the treatment [treatment: F_1,72_=2.6, p>0.05, p*η*^2^=0.03] (**Figure 3D**). Final win-stay scores significantly correlate with final mGT performance [r=0.614, p<0.0000].

At the subpopulation level, good DMs progressively developed and relied on a more rigid and less flexible strategy than intermediate and poor DMs [main effect of cluster, stickiness: F_2,71_=7.0, p<0.01, p*η*^2^=0.16; flexibility: F_2,71_=4.3, p<0.05, p*η*^2^=0.11]; block × cluster interaction, stickiness: F_2,71_=13.5, p<0.0001, p*η*^2^=0.27; flexibility: F_2,71_=9.6, p<0.001, p*η*^2^=0.21]. Intermediate and poor DMs were equally rigid and flexible in their choices [p>0.05, post-hoc] (**Figure S2A, S2B**). Only final stickiness [r=0.551, p<0.0001] and flexibility [r=−0.587, p<0.0000] scores of good DMs significantly correlate with final mGT performance. Concerning the outcome sensitivity, the three DM subpopulations behaved differently along the task [main effect of cluster, lose-shift: F_2,71_=0.7, p>0.05, p*η*^2^=0.02; win-stay: F_2,71_=7.7, p<0.001, p*η*^2^=0.18; block × cluster interaction, lose-shift: F_2,71_= 6.0, p<0.01, p*η*^2^=0.14; win-stay: F_2,71_=7.6, p<0.01, p*η*^2^=0.18]. Initially, good and poor DMs more frequently shift after a penalty than intermediate DMs [p<0.05, post-hoc], the latter significantly increasing their lose-shift-based strategy along the task [p<0.05, post-hoc]. Final lose-shift scores were not different between subpopulations [p>0.05, post-hoc] (**Figure S2C**). Besides, good DMs more frequently chose the same option after a reward as the task progressed [p<0.001, post-hoc], becoming significantly different from intermediate and poor DMs [p<0.01, post-hoc] (**Figure S2D**). Finally, optimal DM relies on final lose-shift [r=−0.447, p<0.001] and win-stay scores [r=0.614, p<0.0000] as they correlate with final mGT performance in good DMs. No effect of the CORT treatment was evidenced for the behavioural dimensions within DM clusters, irrespective of the blocks [good DMs, stickiness, beginning: U=345.5; end: U=336.0; flexibility, beginning: U=302.0; end: U=290.5; lose-shift, beginning: U=325.0; end: U=238.0; win-stay, beginning: U=350.5; end: U=321.5, all ps>0.05; intermediate DMs, stickiness, beginning: U=9.5; end: U=4.5; flexibility, beginning: U=9.0; end: U=8.0; lose-shift, beginning: U=9.0; end: U=10.0; win-stay, beginning: U=9.0; end: U=7.0, all ps>0.05; poor DMs, stickiness, beginning: U=5.5; end: U=3.5; flexibility, beginning: U=2.0; end: U=5.0; lose-shift, beginning: U=2.0; end: U=5.0; win-stay, beginning: U=2.0; end: U=5.0, all ps>0.05].

To better characterize the influence of chronic CORT on DM processes we further addressed its impact on complementary behavioural domains within the RDoC framework. CORT treated mice displayed a more passive coping style when facing uncertainty in the FST, with a significantly longer immobility duration (total time of immobility (s) ± SEM: 175.43±10.36) as compared to SHAM animals (129.07±9.21) [t_76_=−3.3, p<0.01] (**Figure 3E**). As DM subpopulations did not differ [F_2,70_=0.2, p>0.05, p*η*^2^=0.01] (**Figure 3F**) and final mGT performance did not correlate with FST scores [r=−0.12, p>0.05], these results suggest that the coping style does not primarily influence DM performance.

Both SHAM [consumption of sucrose solution vs 50%: t_39_=79.3, p<0.00] and CORT [t_39_=61.6, p<0.00] mice expressed a strong preference for the sucrose solution compared to water (percentage of total sucrose consumption ± SEM, SHAM: 96.50 ± 0.59; CORT: 94.84 ± 0.73) (**Figure 3G**), and no difference between them was evidenced for the sucrose solution consumption [t_78_=1.8, p>0.05]. Moreover, DMs categories did not either differ [F_2,71_=0.7, p>0.05, p*η*^2^=0.02] (**Figure 3H**), suggesting that DM performance does not primarily relies on reactivity to positive outcome.

We further investigated whether chronic CORT alters other dimensions required for goal-directed based DM. Chronic CORT did not impact spatial WM [mean effect of treatment, total number of errors: F_1,41_=1.8, p>0.05, p*η*^2^=0.04; task latency: F_1,41_=3.7, p>0.05, p*η*^2^=0.08]. However, a learning process was highlighted [session, total errors: F_1,41_=4.8, p<0.05, p*η*^2^=0.10; test latency: F_1,41_=5.5, p<0.05, p*η*^2^=0.12] which is accounted for by SHAM animals only [total errors: Z=2.8, p<0.01; test latency: Z=3.3, p<0.001], while CORT mice did not improve along the task [total errors: Z=0.5, p>0.05; test latency: Z=1.2, p>0.05] (**Figures 4A, 4B**). Decision-making subpopulations differed in their WST performance [cluster × session interaction, total number of errors: F_2,40_=7.5, p<0.01, p*η*^2^=0.27; task latency: F_2,40_=5.0, p<0.05, p*η*^2^=0.20], with poor DMs significantly improving through the task [total number of errors: p<0.001; task latency: p<0.01, post-hoc], unlike good [total number of errors: p>0.05; task latency: p>0.05, post-hoc] and intermediate DMs, the latter making more mistakes at the end of the task [total number of errors: p<0.05; task latency: p>0.05, post-hoc]. At the end, only intermediate and poor DMs were different in terms of total number of errors [session 5: F_2,40_=3.4, p<0.05, p*η*^2^=0.14; p<0.01, post-hoc], but not in task latency [session 5: F_2,40_=2.0, p>0.05, p*η*^2^=0.09]. Final WST scores do not correlate with final mGT performance, which questions the influence of spatial WM on DM processes.

**Figure 4.**
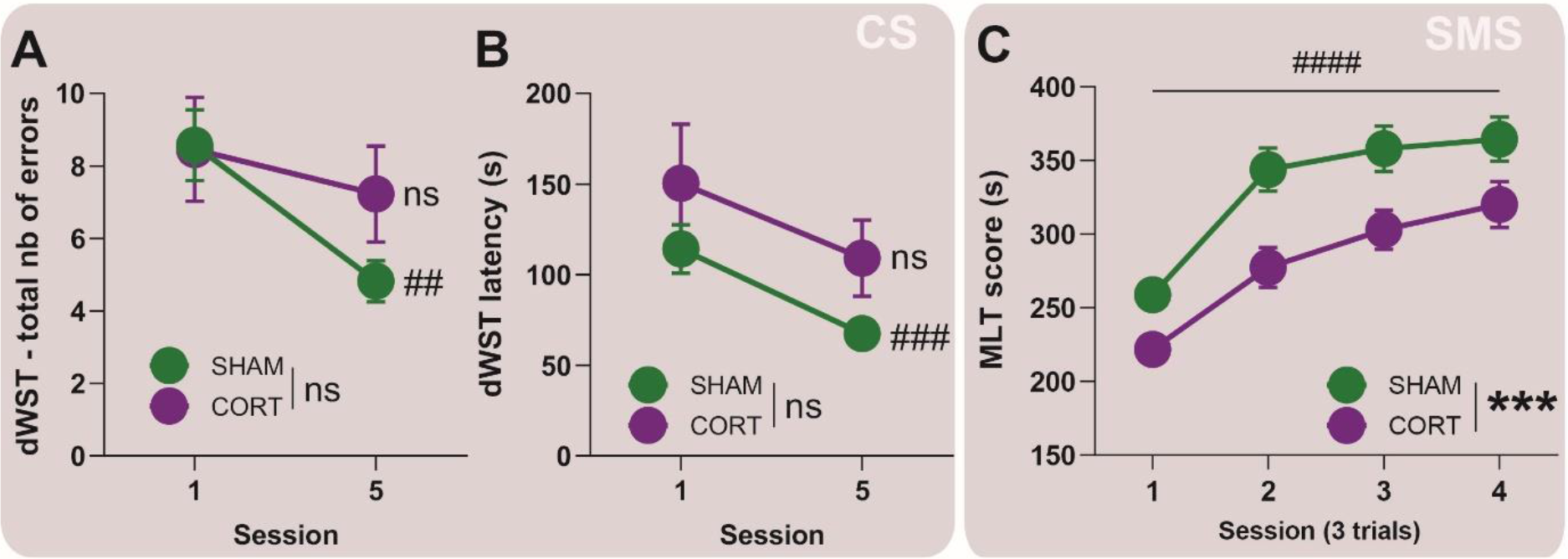
Chronic CORT hampers spatial WM and motor learning processes. Irrespective of the condition [ns], the evaluation of spatial working memory (WM) capabilities in the delayed spatial Win Shift Task (dWST) revealed a general learning process, shown as a decreased in the total number of errors (A) and the time needed to finish the task (B), mainly accounted for by SHAM animals [##, p<0.01; ###, p<0.001] (Cognitive Systems -CS). Additionally, psychomotricity (C) was found to be impaired after chronic CORT exposure [treatment: ***, p<0.001], even if all animals, irrespective of the treatment, improved their performance along the Motor Learning Task (MLT) [session: ####, p<0.0000]. (Sensory Motor Systems -SMS).

As CORT slowdowns onset of optimal DM strategy, we tested whether motor performance sub-serving execution and exploitation of the mGT was also impacted. Mice improved their performance along the task [main effect of session: F_3,636_=38.5, p<0.0000, p*η*^2^=0.15], with chronic CORT impairing their general MLT performance [main effect of treatment: F_1,212_=12.7, p<0.001; p*η*^2^=0.06]. Nevertheless, no significant interaction was found between factors [treatment × session interaction: F_3,636_=0.9, p>0.05, p*η*^2^=0.00]. When comparing individual sessions, CORT animals hold shorter on the rotor than SHAM mice in the first three sessions, a difference that disappeared at the end of the task, thus suggesting a delayed learning process in the pathological condition (**Figure 4C**). Decision-making clusters did not behave differently during the task [main effect of cluster: F_2,211_=0.1, p>0.05; p*η*^2^=0.00] and MLT scores do not correlate with final mGT performance [r=0.0497, p>0.05]. These results show that CORT treatment interferes with motor learning processes, which could impact exploration in the mGT.

We next investigated whether biochemical adaptations to chronic CORT could account for differential DM performance. The FCS appeared significantly degraded in CORT animals from the third week of treatment [main effect of treatment: F_1,890_=448.1, p>0.0000, p*η*^2^=0.33; week: F_13,890_=20.1, p<0.0000, p*η*^2^=0.23; treatment × week interaction: F_13,890_=11.5, p<0.0000, p*η*^2^=0.14; from third week of treatment: all ps<0.01, post-hoc]. DM categories did not differ though, neither in SHAM [main effect of cluster: F_2,36_=1.7, p>0.05, p*η*^2^=0.08] nor in CORT animals [cluster: F_2,32_=0.2, p>0.05, p*η*^2^=0.01]. Moreover, FCS scores do not correlate with final mGT performance [r=0.0205, p>0.05].

As expected, terminal blood sample analyses showed significantly higher basal plasma CORT levels in treated animals (mean CORT level (ng/mL) ± SEM; SHAM: 28.18±4.97; CORT: 79.02±12.41) [t_78_=−3.8, p<0.001], and their HPA axis reactivity to a novel acute stress was significantly blunted (SHAM: 224.02±35.21; CORT: 70.18±29.97) [post-stress plasma CORT levels, U=10.0, p<0.01]. No significant differences in CORT levels were evidenced between DM clusters [main effect of cluster: F_2,71_= 0.3, p>0.05, p*η*^2^=0.01]. CORT levels do not correlate with final mGT performance [r=−0.1491, p>0.05].

Finally, we focused on the key players GR/MR ratio and CRF from the regions of interest. The mPFC GR/MR ratio value of CORT animals (mean ± SEM: 1.55±0.37) was significantly decreased compared to control animals (2.30±0.55) [U=451.0, p<0.05] (**Figure 5B, 5C**). To disentangle the origin of this difference, GR and MR levels were compared separately. While MR levels did not differ between conditions [U=673.0, p>0.05], GR levels were decreased in CORT animals [U=547.0, p<0.05]. No differences were found in the VH, nor the DLS between conditions [effect of treatment, VH: U=447.0; DLS: U=523.0, all ps>0.05]. The three DM clusters do not either differ in their GR/MR ratio, irrespective of the region of interest [cluster, mPFC: H_2,68_=1.3; DLS: H_2,65_=0.3; HV: H_2,61_=0.6, all ps>0.05], suggesting that DM performance do not primarily depend on homeostatic HPA deregulation at the GR level.

**Figure 5.**
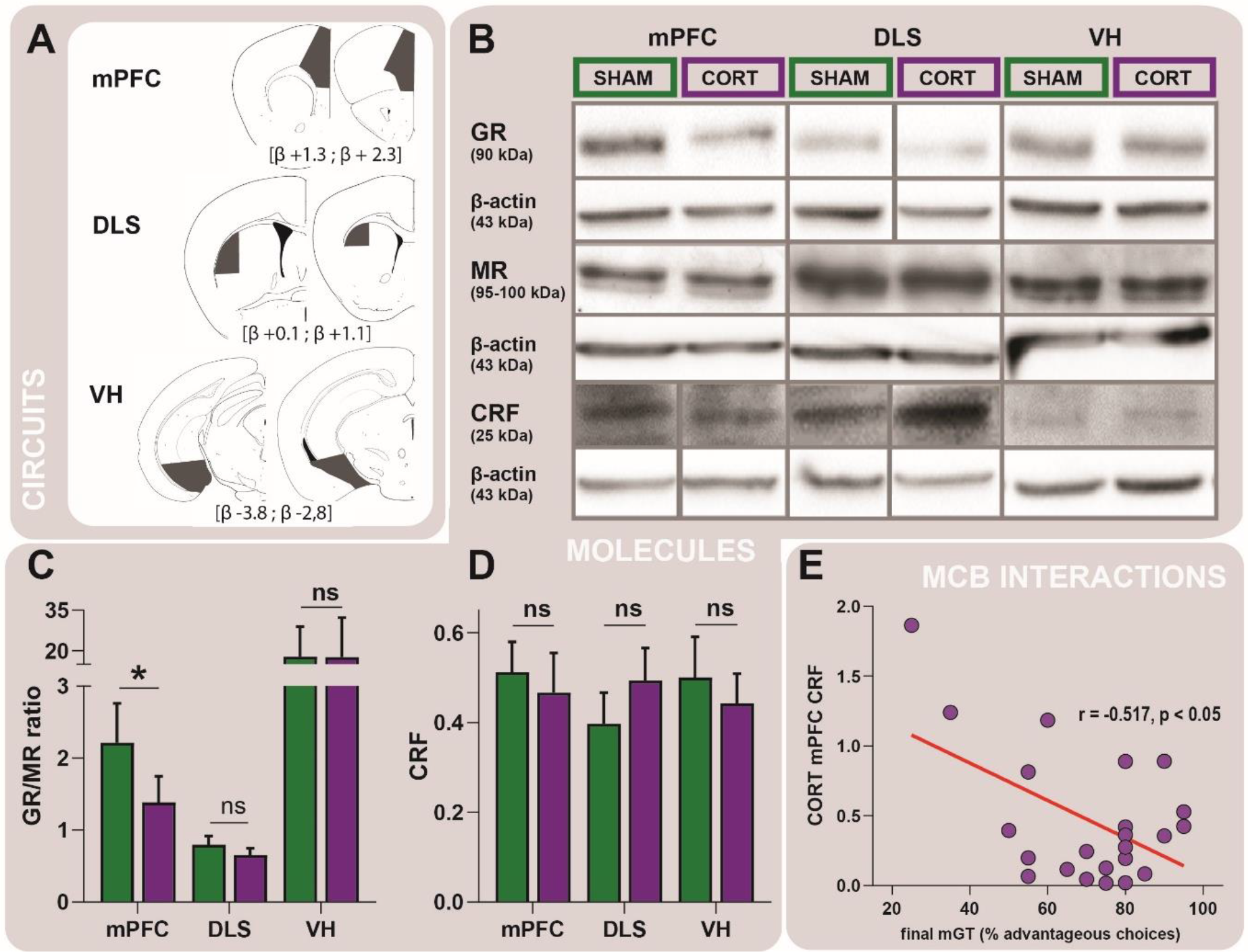
HPA axis imbalance after chronic CORT. (A) Schematic representation of the sampled brain regions’ location for protein quantification. Coronal sections were 1mm-thick. (B) Protein quantification by Western blotting revealed a glucocorticoid receptors (GR) downregulation in the medial prefrontal cortex (mPFC) of CORT-treated animals, which significantly diminished their GR/MR (-mineralocorticoid receptors) ratio value [*, p<0.05] (C), respect to SHAM animals. The protein quantification did not reveal a differential expression in the ventral hippocampus (VH), nor the dorsolateral striatum (DLS), related to the treatment [ns]. (D) Regarding the corticotropin-releasing factor (CRF), chronic CORT did not affect its expression in the brain areas investigated [ns]. However, lower CRF levels in the mPFC correlated with better final mGT performance in CORT [p<0.05] but no SHAM animals. (molecules-circuits-behaviours –MCB-interactions).

Concerning CRF levels and regardless of the brain area, no significant differences between conditions [effect of treatment, mPFC: U=301.0; DLS: U=311.0; HV: U=405.0, all ps>0.05], nor between DM clusters [cluster, mPFC: H_2,49_=1.0; DLS: H_2,52_=0.3; HV: H_2,52_=4.0, all ps>0.05] were evidenced (**Figure 5D**). Nonetheless, a significant correlation between the mPFC CRF levels and the final mGT performance of CORT animals was observed [r=−0.5166, p<0.05] (**Figure 5E**), suggesting that vulnerability to suboptimal DM induced by chronic CORT relates more to CRF signalling deregulation than to GR *per se*.

## DISCUSSION

The results of this study reveal that several weeks of CORT exposure delays the encoding of the contingencies required to select responding towards optimal DM in a probabilistic gambling task. Inter-individual differences in the capability to develop an optimal DM strategy were evidenced in the global mouse population and remarkably, the proportion of individuals displaying suboptimal DM performance is enhanced upon chronic CORT.

The identified chronic CORT-induced suboptimal spatial WM, which seems to be more detrimental to the learning rate than to memory load, could somehow hamper early exploration in the mGT, impending integration of task contingencies, in line with preclinical [53–56] and clinical reports [57]. Besides, chronic CORT, instead of hindering final performance, slows-down the learning rate in the MLT, a DLS-dependent motor task that does not rely on positive valence systems [58], thus evoking clinical psychomotor retardation [59]. Additionally, though reward sensitivity does not directly account for differential DM performance, learning to cope with negative outcomes is impaired upon chronic CORT. Taken together, these results suggest a CORT-induced hedonic misbalance with differential DM performance better accounted for by a dysregulated negative (lose-shift dimension) rather than positive (win-stay dimension and sucrose preference) valence system.

This study disclosed a GR downregulation in the mPFC of treated animals, suggesting that chronic CORT exposure may disturb reflective behaviour for optimal planning, and favour suboptimal habit formation, in line with previous studies addressing the role of the mPFC in instrumental behaviour [39,60,61]. Of particular interest, CRF signalling in the mPFC of CORT individuals was found to negatively correlate with their final DM performance, underlining synergetic effects of CRF and CORT in stress-induced cognitive alterations (in line with [62]).

Experimental studies in humans have proposed that IGT performance relies on reward sensitivity [16] though anhedonia measures do not always correlate with task performance [63]. This is the case for our data in reward sensitivity upon chronic CORT. An efficient exploration phase in the IGT and therefore in its rodent adaptations, would guide the behaviour to a faster stickiness to the optimal choice strategy, that would become more independent of the outcome, i.e. more habitual [39,64]. The spatial WM and psychomotor deficits observed in our CORT-treated mice could affect primarily action-outcome learning crucial for optimal DM strategies, rather than solely cue processing, and even if they do not directly predict differential DM, they may compromise the transition from goal-directed to habit-based behaviour. Taking in consideration the complementary roles of the studied brain structures in instrumental learning [38,39,54], we suggest that the GR downregulation observed in the mPFC of treated animals would primarily entail suboptimal action-outcome encoding over cue processing, yielding action-outcome consolidation especially vulnerable to chronic CORT. This interpretation is in agreement with previous studies reporting negative consequences on cognition upon chronic stress and anxiety [4,65,66].

Stress has been shown to affect *crf* signalling, compromising the positive valence system [67] and disrupting fronto-striatal cognition [20,68]. Together with the present study, these results suggest that high mPFC CRF levels can be considered a neurobiological endophenotype of vulnerability to suboptimal DM under chronic stress. Corticotropin-releasing factor signalling disruption may hinder mPFC computations supporting optimal fronto-striatal cognitive functioning, and specifically goal-directed behaviours, and contribute to overreliance on the negative valence system to form suboptimal action-outcome learning. This interpretation further support the *somatic marker hypothesis* [69,70] and point towards an integral role of CRF. However, previous studies have warned about the dissociable roles of the ventral and dorsal mPFC in DM, especially when behaviour is reward-guided and sensitive to negative feedback [71]. Since we jointly processed ventral and dorsal mPFC structures, further investigations will be necessary to establish the exact contribution of the prelimbic and infralimbic areas of the mPFC in the CRF signalling upon chronic CORT.

The results presented here demonstrate that chronic CORT exposure impedes optimal DM under uncertainty in mice, impacting mPFC GR and CRF signalling. Manipulating the latter to counterbalance overreliance on the negative valence system with suboptimal habit formation, could prove useful to improve coping with risk aversion towards rigidification of optimal choices when DM involves overcoming a conflict.

In sum, this study provides novel insight into the mechanisms of maladaptive value-based DM caused by chronic exposure to GC, and have important implications for understanding pathophysiological mechanisms in a transdiagnostic perspective and for identifying alternative pharmacological targets towards precision medicine in biological psychiatry.

## Acknowledgements

The authors thank Mr Hervé Reyssie (Animal Facilities, Besançon) and Mrs Julie Frejaville (Laboratoire de Carcinogenèse associée aux HPV EA3181, Besançon) for technical support.

## Author contribution

LC, YP and DF conceived the experimental design with input from DB. LC, BR and CH contributed to the data acquisition. LC and YP analysed the data. LC, YP and DB wrote the manuscript. All authors critically revised the work and approved the version to be published. LC, YP and DF agree to be accountable for all aspects of the work in ensuring that questions related to the accuracy or integrity of any part of the work are appropriately investigated and resolved.

**Supplementary Information** accompanies this paper.

