## Supplementary Material for "Chronic Exposure to Glucocorticoids Induces Suboptimal Decision-Making in Mice"

**Chronic CORT Impairs Decision-Making**

MSc Lidia CABEZA[1]*, MSc Bahrie RAMADAN[1], PhD Julie GIUSTINIANI[1,2,3], LA Christophe HOUDAYER[1], PhD Yann PELLEQUER[4], PhD Damien GABRIEL[1,3], PhD Sylvie FAUCONNET[5], Prof Emmanuel HAFFEN[1,2,3], PhD Pierre-Yves RISOLD[1], Prof Dominique FELLMANN[1], PhD David BELIN[6], PhD Yvan PETERSCHMITT[1]*

### SUPPLEMENTARY INFORMATION

#### SUPPLEMENTARY METHODS

##### *Delayed spatial win-shift task (dWST)*

*Individual* spatial working memory (WM) was measured in a delayed spatial WST (dWST) on an 8-arm radial maze over 5 consecutive daily sessions. Mice were habituated to the maze, which consists of 8 identical opaque equidistant arms (37.0 cm long and 5.7 cm wide) and a common central start-point zone (*Panlab,* LE760/2), for three daily sessions prior to dWST training. Three days before the first habituation session, animals were habituated to the reinforcer (20mg Dustless Precision Pellets® Grain-Based Diet, PHYMEP s.a.r.L., Paris, France) by presentation in their home cages. During the first habituation session, animals were placed in the start-point zone and were allowed to explore the maze (all arms open) for 10 minutes. In the second habituation session food pellets were scattered around the maze (all arms open) and animals had 10 minutes two explore and eat. In the last habituation session, one pellet was disposed at the end of each arm, inside a food cup and invisible from the entry of the arm, and animals had again 10 minutes to explore, find and eat the rewards. Mice were then trained for five consecutive daily sessions before the dWST, to visit all 8 arms of the maze. For this, mice were placed on the start-point of the maze, inside an opaque cylindrical structure that prevented visual assessment of the maze by the animal. After 5 seconds the cylinder was removed and all eight arms opened. Mice were allowed to collect one food pellet at the end of each arm and reentries into a previously visited arm were considered an error. The session stopped when the 8 pellets had been collected or after 15 minutes had elapsed. The day following the last training session mice were tested in the dWST for five daily sessions, each of which consisted of a *study* phase and a *test* phase. Briefly, in *study phase* mice were placed on the start-point of the maze, inside the cylindrical structure, and when the cylinder was removed half the arms opened (randomly distributed across mice and sessions) and mice had to collect the available reinforcer in each of the 4 arms. Mice were then brought back to their home cage. After 3 minutes they were placed back on the start-point of the maze for the second phase or *test* phase, during which all 8 arms were opened but only those previously closed during the *study phase* were baited. Reentries in previously visited arms were considered an error. The session lasted until the last pellet was collected, or 15 minutes had elapsed.

##### *Mouse Gambling Task (mGT)*

Decision-making (DM) was measured using the mGT task we have previously described (Cabeza et al., 2020). The task took place in a completely opaque 4-arm radial maze, with identical and equidistant arms, and a common central zone used as a start-point. Mice were rewarded with 20mg grain-based pellets or punished with grain-based pellets previously treated with quinine (180mM quinine hydrocholride, Sigma-Aldrich, Schnelldorf, Germany). Quinine pellets were not palatable but edible.

Mice were trained twice daily for five consecutive days with each daily session consisting of 20 choice trials (a total of 100 trials per animal). On each trial, positive and negative reinforcers were allocated to the arms following a probabilistic rule so that visiting arms *A* and *B* was overall *disadvantageous* while visiting arms C and D was overall *advantageous.* The former resulted in access to immediate larger reward, but larger cumulative negative reinforce in the long run (9 and 14 rewards per session respectively) whereas the latter resulted in access to smaller immediate reward but a smaller cumulative punishment over time (50 and 66 rewards per session). Mice were placed in their home cages for 90 seconds between consecutive trials. The location of advantageous and disadvantageous arms was randomised with different reward and punishment sequences for each animal.

##### *Sucrose preference test (SPT)*

The SPT lasted 4 consecutive nights. On the first night or *forced sucrose phase*, drinking water was substituted by a 2.5% sucrose solution (D(+)Saccharose, Carl Roth, Karlsruhe, Germany) in the home cages. Mice were subsequently given exclusive access over night to two identical bottles, one containing drinking water and the other, the sucrose solution, in individual cages. Every morning bottles were removed and weighted, and animals were placed back in their home cages. The position of the bottles was switched between consecutive sessions in order to avoid side bias. The sucrose preference was calculated as the percentage of sucrose solution consumption, during the last session, relating to the total liquid intake.

##### *Motor learning task (MLT)*

Mice were tested on an accelerating rotarod (3 cm diameter automatized rotor, Mouse RotaRod NG, *Ugo Basile® SRL*, Gemonio, Italy) over 5 consecutive daily sessions. During the first session mice were habituated to stay on the rod with constant speed of 5 rpm for at least 90 seconds. During the 4 following sessions, mice were tested daily over 3 consecutive trials, with an inter-trial interval of 60 seconds. Mice were placed on the rotor with a constant 5 rpm-speed rotation for a minute, and then challenged to stay on the rod despite an acceleration of the motor from 5 to 40 rpm over 5 minutes. Trials ended when the mice fell off the rod or after 10 minutes had elapsed.

##### *Western blotting.*

Individual brain tissue samples were sonicated in lysis buffer RIPA (50 mM Tris-HCl pH 7.4, 150 mM NaCl, 1 mM EDTA, 1% Nonidet P-40, 0.5% sodium desoxycholate) supplemented with protease inhibitors (*Roche Diagnostics*, Meylan, France), centriguged at 10.000 g for 10 minutes at 4°C and stored at -80°C. Protein concentrations were estimated using a Bradford protein assay according to the manufacturer's recommendations (Bio-Rad, Marnes-la-Coquette, France) and 30µg of each sample were solved in Laemmli buffer (Bio-Rad) and separated by a mixed 7.5-12% SDS polyacrylamide gel electrophoresis (GR and MR, and CRF respectively). After migration and transfer to PVDF membranes (*GE Healthcare*, England), blots were saturated with in TBS-Tween 20 buffer (0.5% mM Tris-HCl, 45 mM NaCl, 0.05% Tween 20, pH 7.4) with 5% non-fat milk solution for an hour. Membranes were incubated with primary antibodies overnight at 4°C, and then with conjugated secondary antibodies for 2 hours at room temperature. Membranes were then exposed to chemiluminiscent substrate (ECL Western Blotting Substrate, Pierce™) followed by film exposure (Hyperfilm ECL, *GE Healthcare*) or by using ChemiDoc XRS+ with image lab software (Bio-Rad).

For each antibody, we used protein extracts of different mouse organs: brain, kidney and cerebellum as GR, MR and CRF positive controls respectively: kidney, skeletal muscle and pancreas as negative controls.

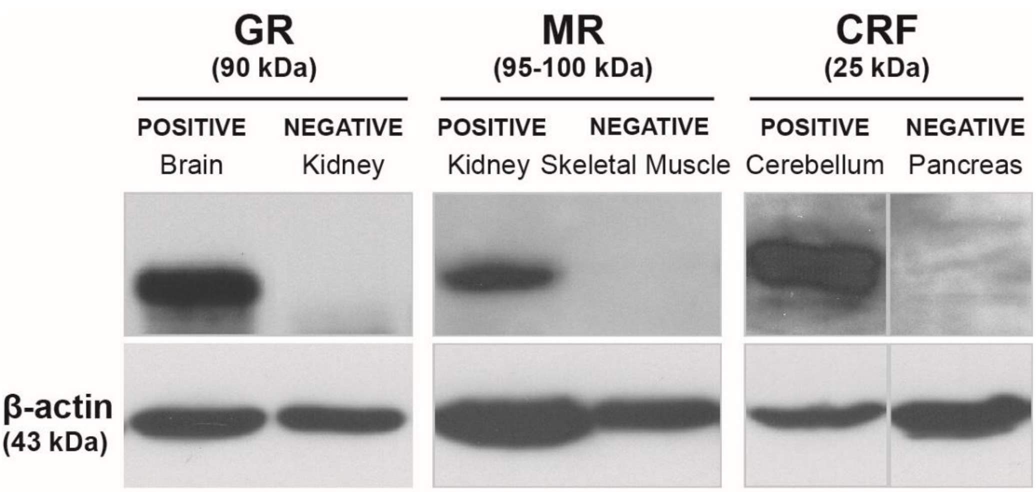


*Figure S1. Western blot positive and negative controls for GR, MR and CRF antibodies.*

#### SUPPLEMENTARY FIGURES

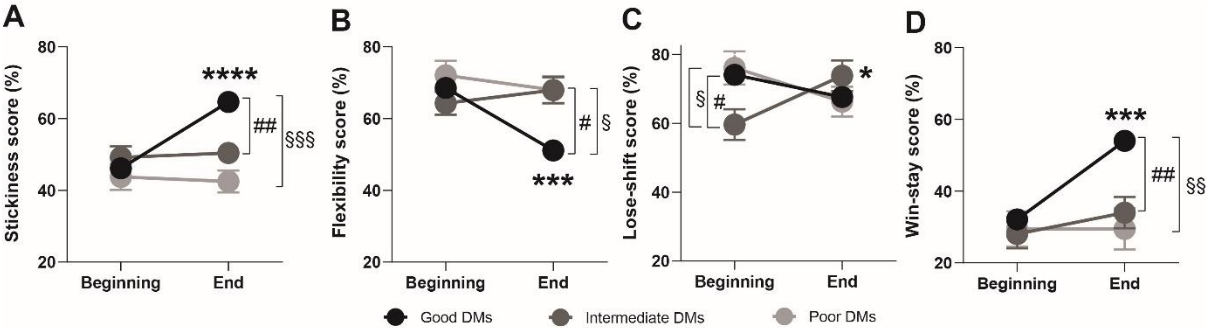


*Figure S2. Characterization of the stickiness, flexibility, lose-shift and win-stay scores of the three DM sub-populations.*

The three DM subpopulations showed pronounced differences with regards to the behavioural dimensions of stickiness [cluster x block interaction: $F_{2,71}$=13.5, p<0.0001] (**A**) and flexibility [$F_{2,71}$=9.6, p<0.001] (**B**). In contrast with that shown by intermediate and poor DM mice, stickiness progressively increased in good DM mice [****, p<0.0001] in parallel with a proportional decrease in flexibility [***, p<0.001], thereby demonstrating the transition from an exploration to exploitation strategy. Thus, by the end of the session, good DM mice showed more stickiness and less flexibility than intermediate DM mice [#, p<0.05; ##, p<0.01], and more stickiness and less flexibility than poor DM mice [§, p<0.05; §§§, p<0.001)], which did not differ from intermediate mice whatsoever. The three subpopulations also differed in their outcome sensibility [cluster x block interaction, lose-shift: $F_{2,71}$= 6.0, p<0.01; win-stay: $F_{2,71}$=7.6, p<0.01]. Intermediate DM mice were less sensitive to the negative outcome at the beginning of the task than good [#, p<0.05] and poor [§, p<0.05] DM mice, and in unlike them, they increased their lose-shift-based strategy along the task [*, p<0.05]. Final lose-shift scores were not different between subpopulations. (**C**). Regarding the sensitivity to a positive outcome, good DM mice more frequently chose the same option after a reward as the task progressed [***, p<0.001], unlike intermediate and poor DM mice. At the end of the task, good DM mice chose significantly more frequently the same option after a reward than intermediate [##, p<0.01] and poor [§§, p<0.01] DM mice.
